# Traditional Norwegian *Kveik* Yeasts: Underexplored Domesticated *Saccharomyces cerevisiae* Yeasts

**DOI:** 10.1101/194969

**Authors:** Richard Preiss, Caroline Tyrawa, George van der Merwe

## Abstract

Human activity has resulted in the domestication of *Saccharomyces cerevisiae* yeasts specifically adapted to beer production. While there is evidence beer yeast domestication was accelerated by industrialization of beer, there also exists a home-brewing culture in western Norway which has passed down yeasts referred to as *kveik* for generations. This practice has resulted in ale yeasts which are typically highly flocculant, phenolic off flavour negative (POF-), and exhibit a high rate of fermentation, similar to previously characterized lineages of domesticated yeast. Additionally, *kveik* yeasts are highly temperature tolerant, likely due to the traditional practice of pitching yeast into warm (>30 °C) wort. Here, we characterize *kveik* yeasts from 9 different Norwegian sources via PCR fingerprinting, phenotypic screens, lab-scale fermentations, and flavour metabolite analysis using HS-SPME-GC-MS. Genetic fingerprinting via interdelta PCR suggests that *kveik* yeasts form a lineage distinct from other domesticated yeasts. Our analyses confirm that *kveik* yeasts display hallmarks of domestication such as loss of 4-vinylguaiacol production and high flocculation, and show superior thermotolerance, ethanol tolerance, fermentation rate, and unique flavour metabolite production profiles in comparison to other ale strains, suggesting a broad industrial potential for this group of yeasts.

## Introduction

It is clear that human activity resulted in the domestication of *Saccharomyces cerevisiae* yeasts specifically adapted for beer production. Recently, it has been shown that present-day industrial beer yeasts have originated from a handful of domesticated ancestors, with one major clade comprising the majority of German, British, and American ale yeasts (“Beer 1” in (Gallone et al., 2016)), and another clade more closely related to wine yeasts which does not have geographic structure (“Beer 2” in (Gallone et al., 2016)). In general, it appears that human selection of beer yeasts over the span of centuries has resulted in the evolution of mechanisms to: efficiently ferment wort sugars such as maltose and maltotriose via duplications of *MAL* genes; eliminate the production of phenolic off flavour (POF) by frequent nonsense mutations in genes, such as *PAD1* and *FDC1*, responsible for production of 4-vinylguaiacol (4-VG) thereby generating POF negative (POF-) strains; and to flocculate efficiently, thereby assisting in the downstream processing of the product (Brown, Murray, & Verstrepen, 2010; Gallone et al., 2016; Gonçalves et al., 2016; McMurrough et al., 1996; Steensels & Verstrepen, 2014).

Regardless of the region of origin, beer yeast was likely maintained and domesticated by reuse (repitching) as well as sharing amongst generations of brewers, resulting in many of the domesticated beer yeasts used in the present day (Gallone et al., 2016; Gibson, Lawrence, Leclaire, Powell, & Smart, 2007; Libkind et al., 2011; Steensels & Snoek, 2014). It must not be assumed, however, that the domestication of beer yeasts occurred solely within the confines of industrial breweries, as there were farmhouse brewing traditions predating the industrialization of beer across northern Europe (Nordland, 1969; Räsänen, 1975). However, the growing industrialization of Europe coupled with convenient commercial yeast availability has abolished traditional home-brewing yeasts in the vast majority of regions, resulting in the loss of regionally unique, domesticated yeasts in the process (Nordland, 1969; Räsänen, 1975; Salomonsson, 1979).

One region where traditional yeast cultures are still being used is western Norway, where a number of farmhouse brewers and/or home brewers have maintained the traditional yeasts of this region, some reportedly for hundreds of years (Fig. 1) (Nordland, 1969). Until recently these yeast cultures, referred to as *kveik*, itself a dialect term for yeast used in this region, were geographically isolated and maintained only locally by traditional farmhouse brewers. It is hypothesized that *kveik* yeasts are domesticated, as beers produced using these yeasts are reported to be non-phenolic (POF-) and the yeasts are reportedly capable of rapidly fermenting malt sugars. Also, much like other domesticated beer yeasts, *kveik* yeasts are maintained and reused via serial repitching (Garshol, 2014; Gibson et al., 2007; Stewart, 2015). However, there are some critical differences in the way *kveik* is used and maintained that may have influenced its adaptive evolution and consequently impacted the generation of specific phenotypic characteristics. First, *kveik* has historically been stored dried on *kveikstokker* (yeast logs) for extended time periods of up to one year or more (Nordland, 1969). Second, *kveik* is typically inoculated with the *kveikstokker* submerged into wort of between 30-40 °C, a very high fermentation temperature for yeast (Caspeta & Nielsen, 2015). Third, this wort is often of high sugar content (up to ~1.080 SG / 19.25 °Plato), and the brewers prefer a short fermentation time, often of only 1-2 days before transferring to a serving vessel (Garshol, 2014; Nordland, 1969). The *kveikstokker* is subsequently withdrawn from the fermentation and dried until its next usage. Taken together, this adaptive environment for *kveik* yeasts was quite different from most ale yeasts, while still favouring the possible development of domesticated traits.

**Fig. 1.**
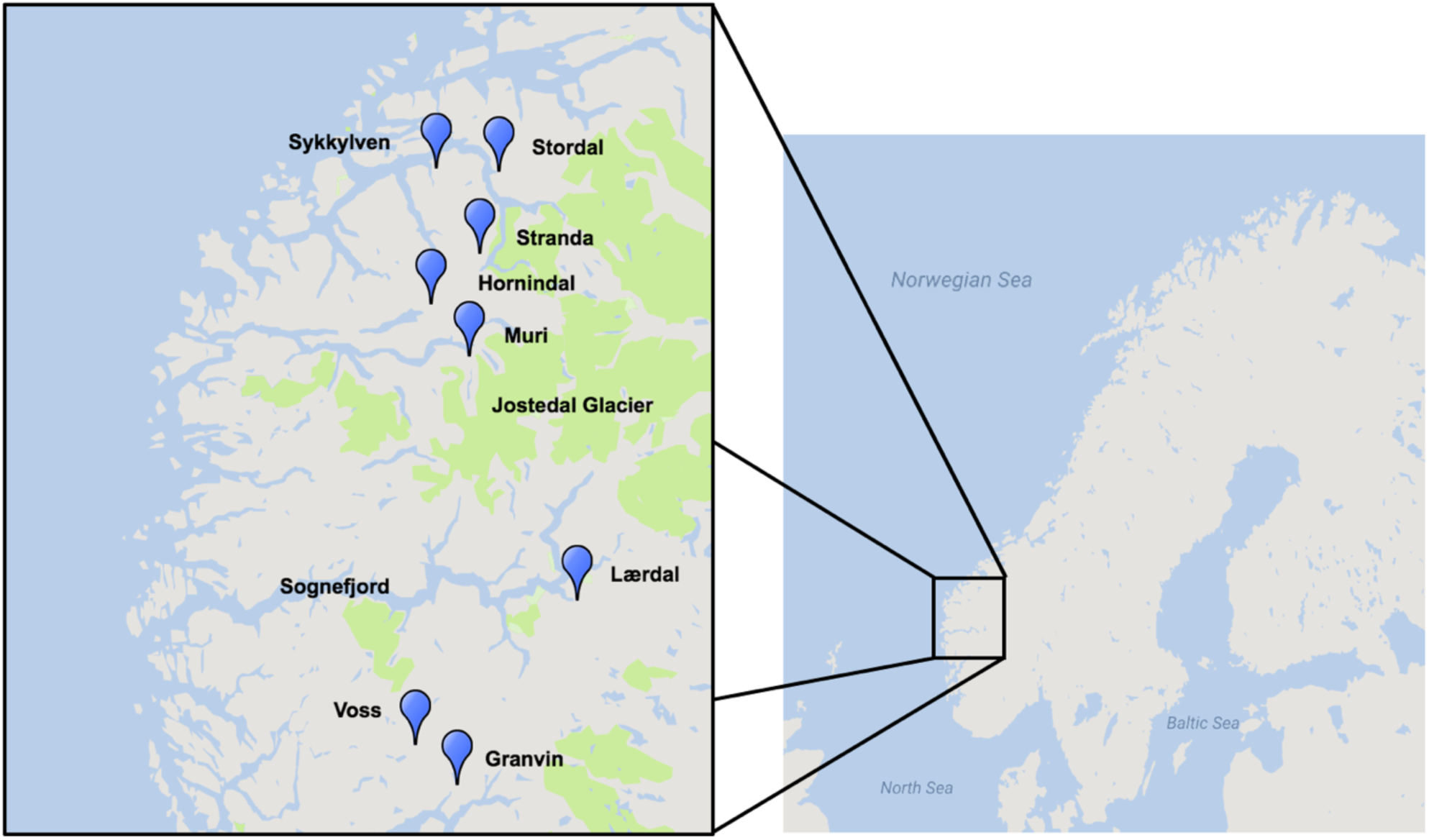
Geographical distribution of *kveik* yeast samples sourced for this project. Map was generated using Google Maps and Scribble Maps. Parks, including the Jostedalsbreen (Jostedal glacier) National Park are highlighted in green.

Remarkably, some *kveikstokker* used for storage of *kveik* can be dated at least as far back as A.D. 1621 (Nordland, 1969), suggesting that *kveik* reuse began well before this date, as presumably the yeast was being reused prior to the development of specialized technology for yeast storage. This lines up with, and potentially predates, recent predictive modelling of the timeline of modern yeast domestication around A.D. 1573-1604 (Gallone et al., 2016). *Kveik* may therefore present a contrasting example of yeasts which have been domesticated and maintained by a geographically isolated brewing tradition, independent of and parallel to industrial beer production.

Yet, critically little is understood about *kveik* yeasts. While some of these yeasts have now been shared globally, there is a lack of cohesive phenotypic and genotypic data pertaining to this intriguing group of beer yeasts. Here we report PCR fingerprinting data that suggest *kveik* yeasts form an interrelated group of ale yeasts genetically distinct from other domesticated ale yeasts. Our phenotypic characterizations reveal classic hallmarks of domestication and, interestingly, unique characteristics in flavour compound production and stress tolerance that increases the potential of *kveik* yeasts in a wide range of industrial applications.

## Materials & Methods

### Yeast Strains

A total of 9 samples of Norwegian *kveik* and one additional Lithuanian farmhouse ale yeast sample were received from a homebrewer in Norway. 7 *kveik* were supplied as liquid slurries, and two were supplied as dried yeast samples. The dried samples were rehydrated in sterile water. The liquid yeast slurries were enriched by inoculating 50 μl of the slurry into 5 mL YPD (1% yeast extract; 2% peptone; 2% dextrose). The samples were incubated at 30 °C for 24 h with shaking, then streak plated onto WLN agar (Thermo Fisher CM0309), which is a differential medium for yeasts, enabling the discrimination of multiple yeasts from one sample. Yeast colonies were then substreaked onto WLN to ensure purity. The resultant strains are summarized in Table 1.

**Table 1.** Investigated yeast strains, source information, and sequence identification. Sequence identification was performed via ITS1-ITS4 rDNA amplification, sequencing, and BLAST.

| Strain Name | Source | Sequence ID | Reference |
| --- | --- | --- | --- |
| Stordal Ebbegarden 1 | Jens Aage Øvrebust; Stordal, Norway | *Saccharomyces cerevisiae* | This study |
| Stordal Ebbegarden 2 | Jens Aage Øvrebust; Stordal, Norway | *Saccharomyces cerevisiae* | This study |
| Stordal Framgarden 1 | Petter B. Øvrebust; Stordal, Norway | *Saccharomyces cerevisiae* | This study |
| Stordal Framgarden 2 | Petter B. Øvrebust; Stordal, Norway | *Saccharomyces cerevisiae* | This study |
| Granvin 1 | Hans Haugse; Granvin, Norway | *Saccharomyces cerevisiae* | This study |
| Granvin 2 | Hans Haugse; Granvin, Norway | *Saccharomyces cerevisiae* | This study |
| Granvin 3 | Hans Haugse; Granvin, Norway | *Saccharomyces cerevisiae* | This study |
| Granvin 4 | Hans Haugse; Granvin, Norway | *Saccharomyces cerevisiae* | This study |
| Granvin 5 | Hans Haugse; Granvin, Norway | *Saccharomyces cerevisiae* | This study |
| Granvin 6 | Hans Haugse; Granvin, Norway | *Saccharomyces cerevisiae* | This study |
| Granvin 7 | Hans Haugse; Granvin, Norway | *Saccharomyces cerevisiae* | This study |
| Granvin 8 | Hans Haugse; Granvin, Norway | *Saccharomyces cerevisiae* | This study |
| Granvin 9 | Hans Haugse; Granvin, Norway | *Saccharomyces cerevisiae* | This study |
| Hornindal 1 | Terje Raftevoll; Hornindal, Norway | *Saccharomyces cerevisiae* | This study |
| Hornindal 2 | Terje Raftevoll; Hornindal, Norway | *Saccharomyces cerevisiae* | This study |
| Hornindal 3 | Terje Raftevoll; Hornindal, Norway | *Saccharomyces cerevisiae* | This study |
| Joniškėlis | Julius Simonaitis; Joniškėlis, Lithuania | *Saccharomyces cerevisiae* | This study |
| Lærdal 1 | Dagfinn Wendelbo; Lærdal, Norway | *Saccharomyces cerevisiae* | This study |
| Lærdal 2 | Dagfinn Wendelbo; Lærdal, Norway | *Saccharomyces cerevisiae* | This study |
| Muri | Bjarne Muri; Olden, Norway | *Saccharomyces cerevisiae* / *eubayanus* / *uvarum* | This study |
| Stranda | Stein Langlo; Stranda, Norway | *Saccharomyces cerevisiae* | This study |
| Sykkylven 1 | Sigurd Johan Saure; Sykkylven, Norway | *Saccharomyces cerevisiae* | This study |
| Sykkylven 2 | Sigurd Johan Saure; Sykkylven, Norway | *Saccharomyces cerevisiae* | This study |
| Voss 1 | Sigmund Gjernes; Voss, Norway | *Saccharomyces cerevisiae* | This study |
| Voss 2 | Sigmund Gjernes; Voss, Norway | *Saccharomyces cerevisiae* | This study |
| WLP001 | USA; White Labs | *Saccharomyces cerevisiae* | This study, (Rogers, Veatch, Covey, Staton, & Bochman, 2016) |
| WLP002 | USA; White Labs | *Saccharomyces cerevisiae* | This study |
| WLP029 | USA; White Labs | *Saccharomyces cerevisiae* | This study |
| WLP570 | USA; White Labs | *Saccharomyces cerevisiae* | This study, (Kopecká, Němec, & Matoulková, 2016) |
| WLP090 | USA; White Labs | *Saccharomyces cerevisiae* | This study |
| WLP007 | USA; White Labs | *Saccharomyces cerevisiae* | This study, (Kopecká et al., 2016) |
| BBY002 | Canada; Escarpment Laboratories | *Saccharomyces cerevisiae* | This study |
| WY1272 | USA; Wyeast | *Saccharomyces cerevisiae* | This study |
| WY1007 | USA; Wyeast | *Saccharomyces cerevisiae* | This study |
| WY2565 | USA; Wyeast | *Saccharomyces cerevisiae* | This study |
| WY1318 | USA; Wyeast | *Saccharomyces cerevisiae* | This study |

### DNA Extraction

DNA was extracted using an adaptation of a previously described method (Ausubel et al., 2002). Briefly, yeast cells were grown in 3 mL of YPD broth at 30 °C, 170 rpm for 24 h, washed with sterile water, and pelleted. The cells were resuspended in 200 μL of breaking buffer (2% Triton X-100, 1% SDS, 100 mM NaCl, 10 mM Tris-HCl). 0.3 g of glass beads and 200 μL of phenol/chloroform/isoamyl alcohol was added and the samples were vortexed continuously at maximum speed for 3 min to lyse the cells. Following centrifugation, the aqueous layer was transferred to a clean tube and 1 mL of 100% ethanol was added. The supernatant was removed following another centrifugation step. The resulting pellet was resuspended in 400 μL of 1X TE buffer and 30 μL of 1 mg/mL DNase-free RNase A and incubated at 37 °C for 5 min. The pellet was then washed with 1 mL of 100% ethanol and 10 μL of 4 M ammonium acetate, followed by another wash with 1 mL of 70% ethanol, and then resuspended in 100 μL of sterile ddH_2_O.

### PCR and Sequencing

The internally transcribed spacer (ITS) regions of the yeast strains were amplified using ITS1 and ITS4 primers (Pham et al., 2011). PCR reactions contained 1μL of genomic DNA, 2.5 μM of each primer, 0.4 mM dNTPs, 2.5 U of Taq DNA polymerase, and 1X Taq reaction buffer. The amplification reactions were carried out in a BioRad T100 Thermocycler under previously described conditions (Pham et al., 2011). PCR products were visualized on a 1% agarose gel in 1X TAE buffer to ensure successful amplification. The samples were purified using the QIAquick PCR purification kit and sequenced using an Applied Biosystems 3730 DNA analyzer. 4peaks software was used to perform quality control of sequence traces. The resulting sequences were analyzed for species-level homology using NCBI BLAST (blastn suite).

### DNA Fingerprinting

Yeast strains were identified by interdelta PCR fingerprinting using interdelta primers *δ*2(5’-GTGGATTTTTATTCCAACA-3’), *δ*12 (5’-TCAACAATGGAATCCCAAC-3’) and *δ*21 (5’-CATCTTAACACCGTATATGA-3’) (Legras & Karst, 2003; Ness, Lavallee, Dubourdieu, Aigle, & Dulaub, 1993). Primer pairs selected for further amplification and analysis were *δ*2+*δ*12 and *δ*12+*δ*21, which both yielded the greatest range of well-resolved bands. PCR was carried out as follows: 4 min at 95 °C, then 35 cycles of 30 s at 95 °C, 30 s at 46 °C, then 90 s at 72 °C, followed by a final 10 min step at 72 °C (Legras & Karst, 2003). Reaction products were confirmed through electrophoresis on a 1% agarose gel in 1X TAE buffer. PCR samples were then purified using a QIAquick PCR purification kit and analyzed on an Agilent 2100 Bioanalyzer using the Agilent DNA 7500 chip. Banding patterns obtained using Bioanalyzer were analyzed using GelJ software. Comparisons for each primer set (*δ*2+*δ*12 and *δ*12+*δ*21) were generated independently using the Comparison feature of the software, clustering the fingerprints using Pearson correlation and UPGMA (Heras et al., 2015). Resultant individual distance matrices were combined using fuse.plot in R (https://github.com/andrewfletch/fuse.plot), which uses the hclust algorithm to format and fuse the matrices and perform hierarchical clustering with UPGMA. The data were visualized using FigTree software (http://tree.bio.ed.ac.uk/software/figtree/).

### Wort Preparation

Wort used for beer fermentations and yeast propagation was obtained from a commercial brewery, Royal City Brewing. The hopped wort was prepared using 100% Canadian 2-row malt to an original gravity of 12.5 °Plato (1.050 specific gravity). The wort was sterilized prior to use at 121 °C for 20 min, and cooled to the desired fermentation or propagation temperature overnight.

### Propagation and Fermentation

Colonies from WLN plates were inoculated into 5 mL of YPD and grown at 30 °C, 170 rpm for 24 h. The YPD cultures were transferred into 50 mL of sterilized wort and grown at 30 °C, 170 rpm for 24 h. These cultures were counted using a haemocytometer and inoculated at a rate of 1.2x10^7^ cells/mL into 70 mL of sterilized wort in glass ‘spice jars’ fitted with airlocks. These small-scale fermentations were performed in triplicate at 30 °C for 12 days. The jars were incubated without shaking to best approximate typical beer fermentation conditions. Fermentation profiles were acquired by weighing the spice jars to measure CO_2_ release, normalizing against water evaporation from the airlocks.

### Beer Metabolite Analysis

Following fermentation, samples were collected and filtered with 0.45 μm syringe filters prior to metabolite analysis. Flavour metabolite analysis was performed using HS-SPME-GC-MS, with a method adapted from Rodriguez-Bencomo *et al.* (2012) (Rodriguez-Bencomo et al., 2012). Samples contained 2 mL of beer, 0.6 g of NaCl, 10 μL of 3-octanol (0.01 mg/mL), and 10 μL of 3,4-dimethylphenol (0.4 mg/mL). 3-octanol and 3,4-dimethylphenol were used as internal standards. The ethanol content was measured using HPLC and a refractive index (RI) detector. The samples were analyzed using an Aminex HPX-87H column, using 5 mM sulfuric acid as the mobile phase, under the following conditions: flow rate of 0.6 mL/min, 620 psi, and 60 °C. Each sample contained 400 μL of filtered beer and 50 μL of 6% (v/v) isopropanol as the internal standard.

### Phenotypic Assays

To determine temperature tolerance, yeast cultures grown for 24 h at 170 rpm at 30 °C in YPD were subcultured into YPD pre-warmed to specified temperatures (15 °C, 30 °C, 35 °C, 40 °C, 42 °C, 45 °C) and grown with shaking for 24 h at the indicated temperature (48 h in the case of 15 °C) and assessed visually for growth. Typical growth (as compared to the same strain at 30 °C) was scored as '+', while reduced growth was scored as '+/-'. No growth was scored as “-“.

To determine ethanol tolerance, yeast cultures grown for 24 h at 170 rpm at 30 °C in YPD were sub-cultured into YPD with increasing concentrations of ethanol (YPD + EtOH 5%; 7%; 9%; 11%; 12%; 13%; 14%; 15%; 16%) to an initial density of 0.1 OD_600_ in 200 μL media in sterile 96 well plates, and grown with shaking at 30 °C for 24 h before being assessed visually for growth. Unimpaired growth (as compared to 5% ethanol) was scored as '+', while reduced but visible growth was scored as '+/-'. No growth was scored as “-“.

To determine flocculation, yeast cultures were grown for 24 h at 170 rpm at 30 °C in YPD, and then 0.5 mL was inoculated into 5 mL sterilized wort, which was incubated for 24 h at 170 rpm at 30 °C. Flocculation was assessed using the spectrophotometric absorbance methodology of ASBC method Yeast-11 (ASBC, n.d.). Values are expressed as % flocculance, with <20% representing non-flocculant yeast and >85% representing highly flocculant yeast.

## Results

### *Kveik* yeasts are genetically distinct from other groups of domesticated yeasts

Given anecdotal reports that *kveik* cultures harbour multiple yeast strains, the *kveik* samples were first plated on WLN agar, which is a differential medium allowing for distinguishing of *Saccharomyces* on the basis of differences in colony morphology and uptake in the bromocresol green dye (Hutzler et al., 2015). Indeed, we found that all but two of the *kveik* samples contained more than one distinct yeast colony morphology, corresponding to hypothetically unique strains (Table 1). The number of putative strains isolated from the *kveik* cultures thus ranged from 1 to 8.

Given that anecdotal reports stated *kveik* yeasts are often flocculent, demonstrate a fast fermentation rate, and are capable of utilizing malt sugars, all of which are hallmarks of domestication (Gallone et al., 2016), we sought to determine whether these yeasts may represent a separate family among previously established families of domesticated ale yeasts, or whether they formed an admixture among existing ale yeast groups. As nearly all domesticated yeasts belong to the *Saccharomyces cerevisiae* species, we hypothesized that the *kveik* isolates also belong to *Saccharomyecs cerevisiae* (Almeida et al., 2015; Gallone et al., 2016; Gonçalves et al., 2016). We performed ITS sequencing and found that all but one *kveik* strain was identified (via BLAST search) as *S. cerevisiae* (Table 1). We found that the strain originating from Muri is most closely homologous to previously identified *S. cerevisiae / eubayanus / uvarum* triple hybrids, presenting this particular yeast strain as an intriguing potential domesticated hybrid warranting further investigation (Table 1).

Since the *kveik* yeasts appear to be *S. cevevisiae* strains, we next asked how they relate genetically to other domesticated ale yeasts. In order to answer this question, we performed interdelta PCR, an example of amplified fragment length polymorphism (AFLP-PCR) using the *δ*12/21 and *δ*2/12 primer sets (Hutzler et al., 2015; Legras & Karst, 2003). The 8 elements are separated by amplifiable distances in the *Saccharomyces cerevisiae* genome, and consequently interdelta PCR can be used to amplify interdelta regions, which in turn can be used to rapidly fingerprint yeasts for comparative genetic purposes (Hutzler et al., 2015; Legras & Karst, 2003).

Preliminary trials using the *δ*1/2, *δ*2/12, and *δ*12/21 primer sets showed that the latter two primer sets produced the greatest range of useful bands when separated via agarose gel electrophoresis. We then amplified the *δ*2/12 and *δ*12/21 regions of all the *kveik* strains and a selection of other ale yeast strains. Separation was performed using capillary gel electrophoresis (Agilent Bioanalyzer), which yielded greater accuracy and sensitivity (Hutzler et al., 2015). Analysis of both *δ*2/12 and *δ*12/21 datasets individually revealed that the *kveik* yeasts formed a subgroup among the other ale yeasts, such that the *kveik* yeasts appeared to be more closely related to each other than to other domesticated ale yeasts (Fig. S1). We next created a composite analysis of the interdelta datasets, yielding a dendrogram which closely matched the expected families of domesticated ale yeasts as previously determined (Fig. 2) (Gallone et al., 2016). We found that the English, American, and German ale families were represented in the dendrogram, but that the *kveik* yeasts mostly did not fit into any of these families. Instead, *kveik* formed a separate group on the dendrogram distantly related to the other groups of domesticated yeasts, with the most closely related group appearing to be a clade containing two German ale strains and one British strain (Fig. 2).

**Fig. 2.**
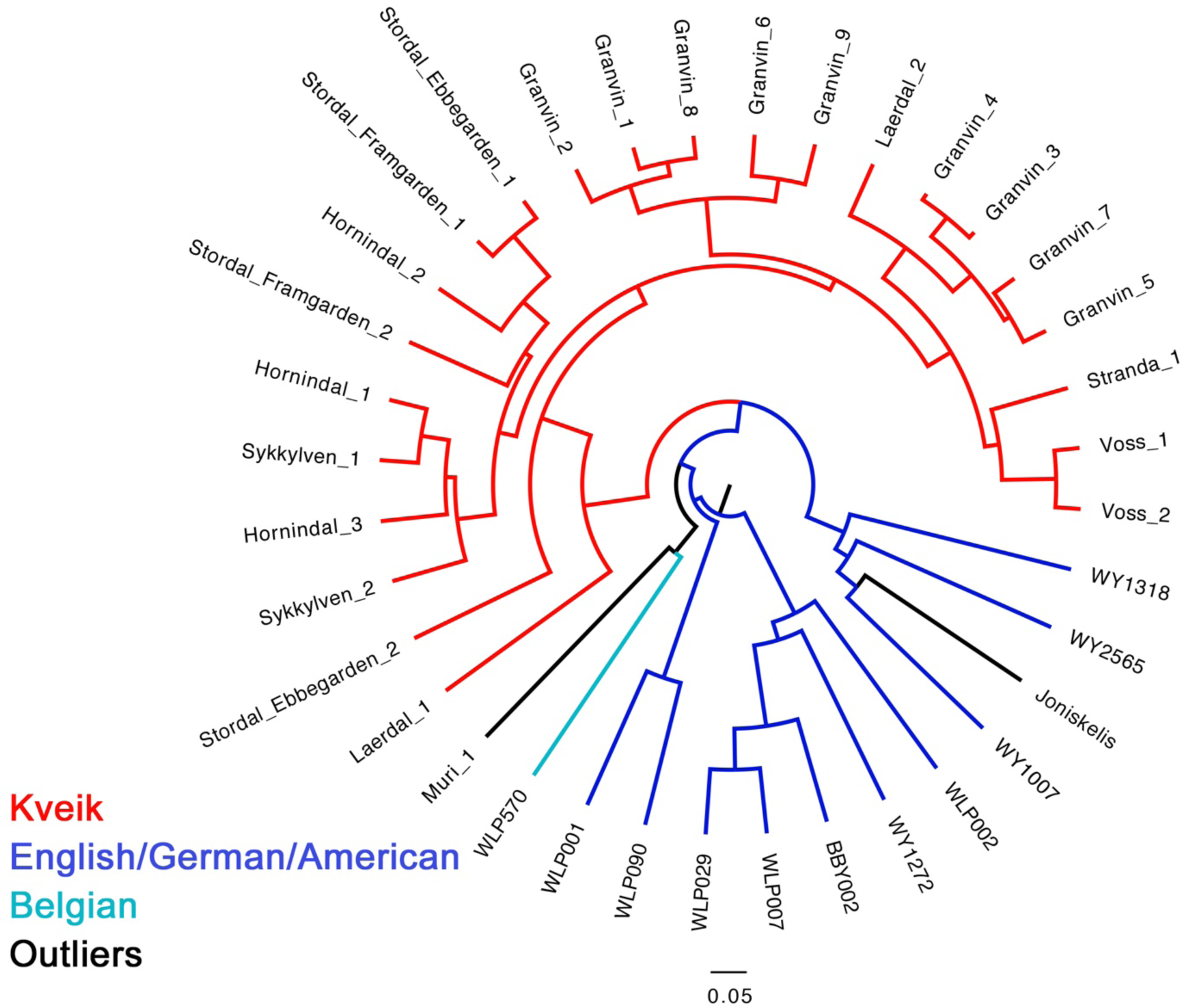
Genetic relatedness of kveik yeasts and other ale yeasts constructed via composite analysis of *δ*12/21 and *δ*2/12 interdelta PCR fingerprints. Composite data were obtained by analysis of individual primer sets in GelJ software, generating difference tables using Pearson correlation and UPGMA. Difference tables were merged using hclust hierarchical clustering algorithm in R. Tree data (Newick) was exported and visualized using Figtree software.

Interestingly, we found that among the *kveik* yeasts, there was also a separation of two major groups, one consisting of strains from Granvin, Stranda, Laerdal and Voss origin, and the other from Sykkylven, Hornindal, and Stordal. It is worth noting that with exception to the Stranda strain, these *kveik* groups correspond geographically to north (Sykkylven, Hornindal, Stordal) and south (Granvin, Laerdal, Voss) of the Jostedal glacier, a major geographic and cultural divide in this region of western Norway (Fig. 1). Thus it appears that the investigated *kveik* yeasts can be divided into two main families with geographic structure on the basis of our fingerprint analysis (Fig. 1, 2). Further, Laerdal and Stordal strains (Laerdal I, Stordal Ebbegarden II) appear within the *kveik* family but more distantly related to the two major groups. Additionally, other yeasts from this study such as the hybrid Muri yeast and the Lithuanian yeast strain do not fit within the *kveik* family. Nonetheless, our data indicate that the majority of *kveik* yeasts represent a new branch on the ale yeast family tree distinct from the genomically characterized English, American, and German ale yeast groups (insert reference for genomic characterization).

### *Kveik* yeasts show strong positive brewing characteristics

We next sought to analyze the brewing-relevant parameters of *kveik* yeasts in pure culture fermentation. Since Norwegian *kveik* cultures appear to often contain multiple yeast strains, there is the possibility that strains are interdependent. It is therefore important to determine the fermentation characteristics of individual strains as single culture fermentations would show whether individual *kveik* strains can adequately ferment beer. An inability to do so would suggest there is an adapted advantage to the multi-strain nature of *kveik* cultures. Additionally, we aimed to confirm anecdotal reports that these yeasts exhibit extremely short lag phases and display good fermentation kinetics.

We performed test fermentations using the pure culture *kveik* strains and an American ale yeast (WLP001; White Labs) as a control, at 30 °C which has been reported to be a typical temperature for beers fermented using *kveik* (Garshol, 2015). In order to assess the fermentation rate during the early phases of wort fermentation, we monitored the CO_2_ loss in the fermentations via weighing. Using this technique, we found that 19/25 *kveik* strains outperformed the American ale control at 24 h, with the best-performing strain producing 91% more CO_2_ within the first 24 h of fermentation (Fig. 3A). Following the 12-day fermentation and maturation period, we also measured ethanol concentration of the beers using HPLC. We found that the *kveik* yeasts produced expected ethanol yields for beer strains of *S. cerevisiae*, with apparent attenuation ranges spanning 65-95%, and ethanol yield ranging from 4.43% ± 0.35% to 6.44% ± 0.46% (Fig. 3B). Furthermore, 15/25 strains produced ethanol values within ± 0.5% of the American ale control, indicating that *kveik* yeasts attenuate wort within the expected range of industrial domesticated ale yeasts. This data confirms that *kveik* yeasts are domesticated as maltotriose in the wort has been utilized or partially utilized, suggesting that *kveik* may also have undergone duplication of *MAL* genes like other domesticated yeasts (Gallone et al., 2016). Furthermore, it appears that *kveik* strain selection, much like selection of individual strains of other domesticated yeasts, is a viable tool to target precise attenuation degrees for targeted beer sweetness/dryness.

**Fig. 3.**
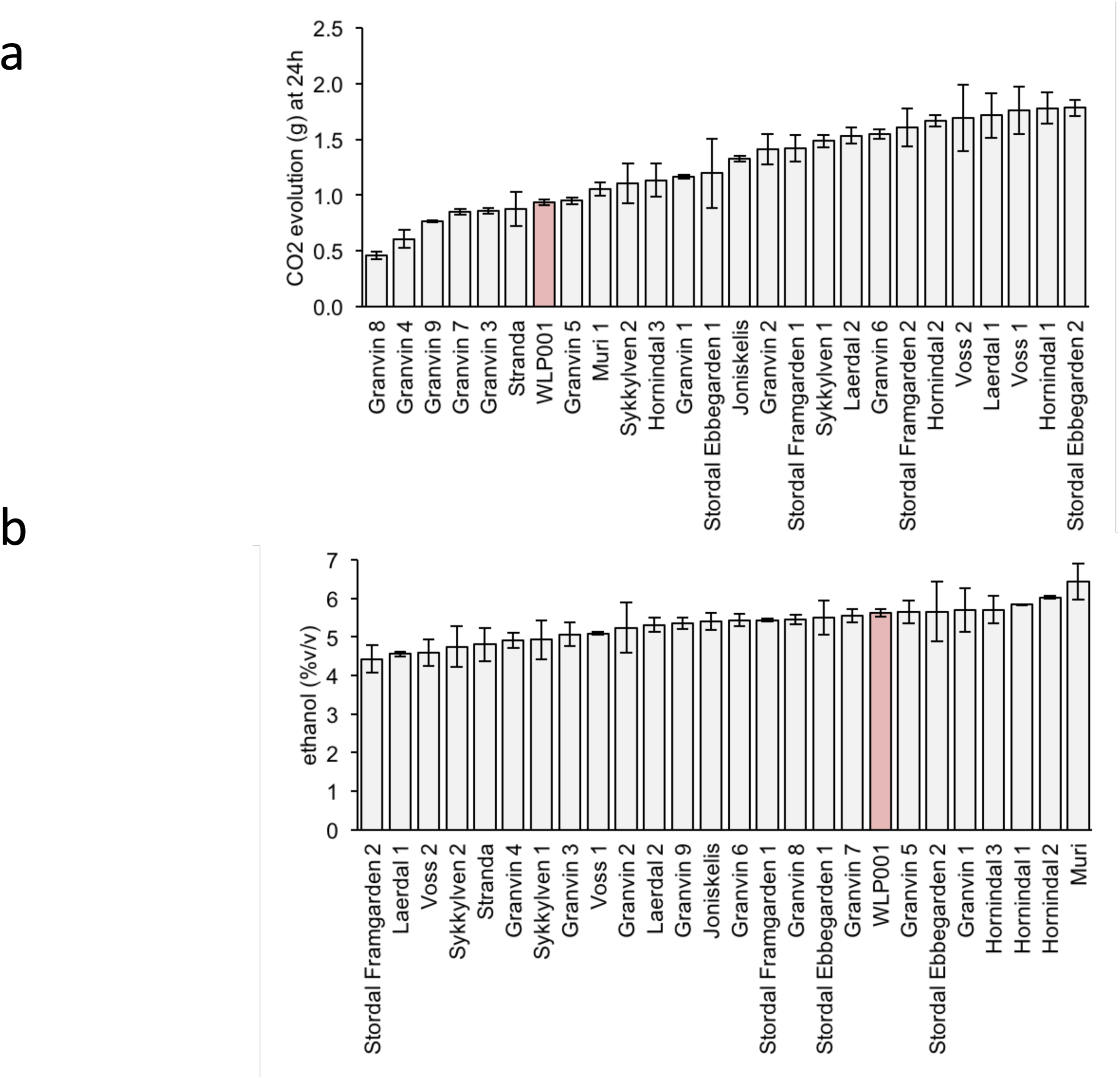
Fermentation kinetics and terminal ethanol concentration of wort fermentation (12.5°P original density) at 30 °C. a) CO_2_ evolution at 24 h was calculated by weighing the fermentation vessels (70 mL) and normalizing for weight loss in the fermentation airlocks. Error bars represent SEM, n=3. Control ale strain is marked in light red. b) Ethanol concentration was measured via HPLC following 12 days of fermentation. Error bars represent SEM, n=3. Control ale strain is marked in light red.

To determine flavour contributions by the *kveik* yeasts, we also analyzed volatile aromatic compounds using HS-SPME-GC-MS (Table 2). Intriguingly, we found that all *kveik* yeasts belonging to the main *kveik* genetic lineage (Fig. 2) produced minimal levels of 4-vinylguaiacol, indicating that the *kveik* family are POF-similar to other domesticated yeast families that has lost the functions of its *PAD1* and *FDC1* genes (Table 2) (Gallone et al., 2016; Gonçalves et al., 2016). Also, analysis of the volatile ester profiles revealed the *kveik* yeasts produced above-threshold concentrations of three yeast fatty acid esters: ethyl caproate (pineapple, tropical), ethyl caprylate (tropical, apple, cognac), and ethyl decanoate (apple) (Comuzzo, Tat, Tonizzo, & Battistutta, 2006; Verstrepen et al., 2003). One or more of these esters was present at above-threshold levels in all of the *kveik* yeasts. Phenethyl acetate (honey, floral, yeasty) was also detected at above-threshold level in 5/25 *kveik* strains. These data suggest that *kveik* yeasts present a potential new option as POF-ale yeasts with a range of intensities of desirable fruity esters.

**Table 2.**
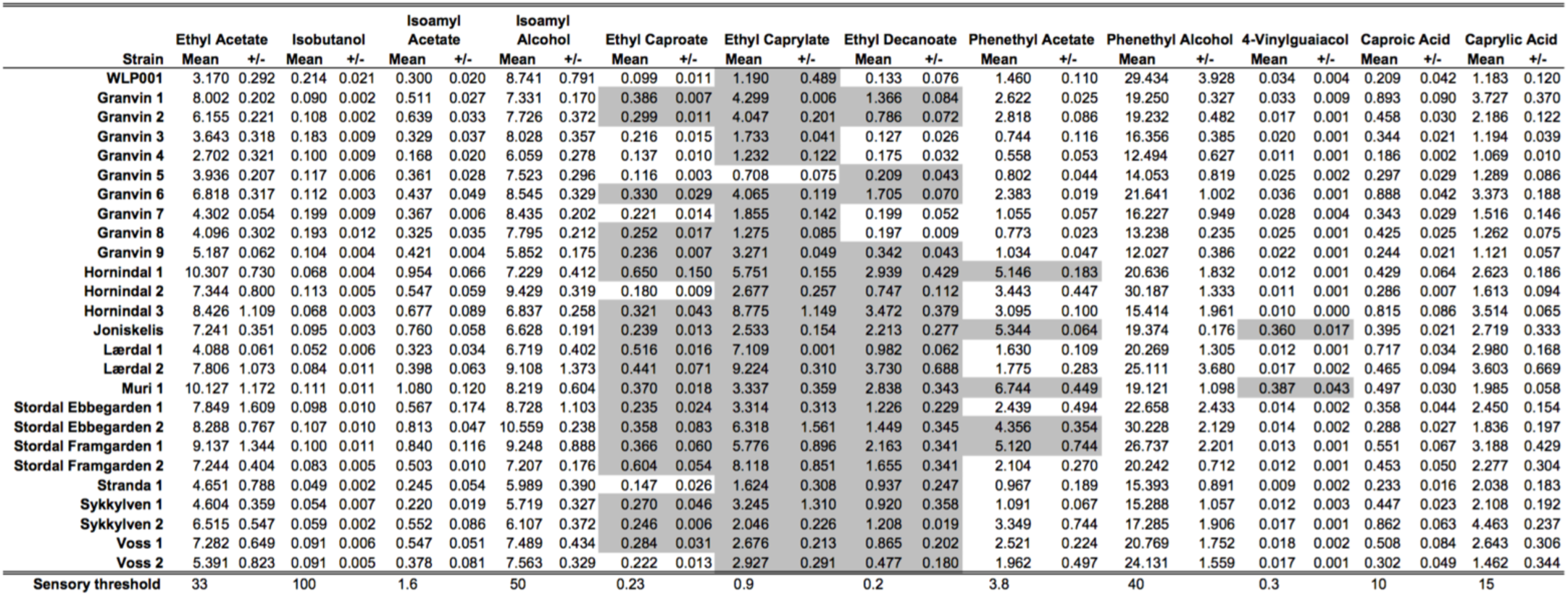
Fermentation flavour metabolites produced by kveik yeasts during wort fermentation (12.5 °P) at 30 °C measured using HS-SPME-GC-MS. Fermentations were performed in triplicate. Metabolites determined to be above the sensory threshold for a given compound are shaded in grey. Values presented are in ppm.

### *Kveik* yeasts demonstrate superior thermotolerance, ethanol tolerance, and flocculation

Since the initial fermentation trials demonstrated *kveik* yeasts are largely POF-and produce desirable fruity ester flavours, we next investigated the stress tolerance and flocculation of these yeasts to better determine their potential utility and to confirm these additional hallmarks of domestication. We monitored the growth of the *kveik* yeasts alongside known ale yeast control strains (WLP001; American ale, WLP029; German ale, WLP570; Belgian ale, WLP002; British ale) over a broad range of temperatures (15 °C to 45 °C), given the reports of high-temperature fermentation by traditional Norwegian brewers (Garshol, 2014; Nordland, 1969). We found that while the control commercial yeast strains did not grow above 40 °C, and several were limited to lower temperatures, all *kveik* strains grew at 40 °C (Table 3). Remarkably, 18/24 *kveik* strains displayed uninhibited or reduced but visible growth at 42 °C, nearing the theoretical limit, and current technological upper threshold for *S. cerevisiae* cell growth (Caspeta et al., 2013; Caspeta, Chen, & Nielsen, 2016; Caspeta & Nielsen, 2015). Interestingly, two from Granvin and two from Voss, showed entirely uninhibited growth at 42 °C. Thus, it appears that thermotolerance is a conserved feature among *kveik* yeasts. However, thermotolerance as a domestication marker has not been fully explored, but may be a distinguishing adaptation of the *kveik* family of yeasts.

**Table 3.**
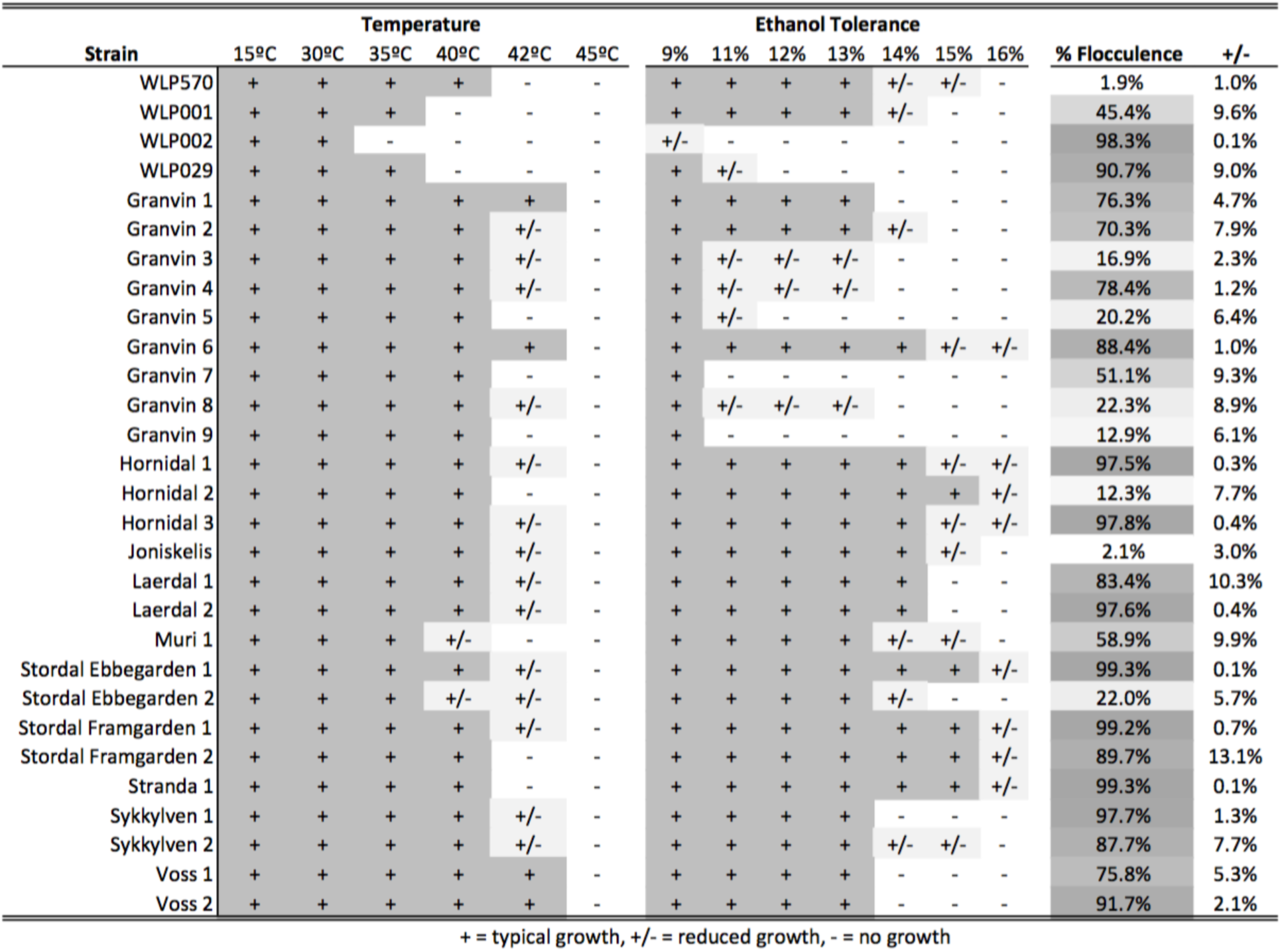
*Kveik* yeasts demonstrate high levels of thermotolerance, ethanol tolerance, and flocculation. Temperature tolerance, ethanol tolerance, and flocculence assays were performed as described in materials and methods. For temperature and ethanol tolerance assays, boxes showing typical growth are shaded darkly and reduced growth is partially shaded. For flocculence assay, boxes are shaded according to degree of flocculence, with darker shading indicating higher flocculation.

We next investigated the ethanol tolerance of *kveik* yeasts in comparison to commercial ale strains with ethanol tolerances available from the suppliers (White Labs, Wyeast, Escarpment Laboratories). Our control data was in line with the suppliers’ broadly specified ethanol tolerances, eg. WLP001 to ‘High – 10-15%’ (13%; Table 3) and WLP002 to ‘Medium – 5-10%’ (9%; Table 3). For the *kveik* yeasts, we found that 15/25 strains displayed typical growth or reduced growth at 14% ethanol, and likewise 11/25 strains for 15% ethanol. 8/25 strains grew in 16% ethanol; however, all samples showed reduced growth at this ethanol concentration (Table 3). With exception to a number of strains originating from the Granvin sample, *kveik* yeasts display high levels of ethanol tolerance, providing further support of domestication of these yeasts.

Flocculation is a hallmark of yeast domestication, as this property enhances the brewer’s ability to harvest yeast via either top or bottom cropping in the fermentor (ref). We assessed the flocculence of the *kveik* yeasts using the absorbance method of ASBC Yeast-11 Flocculence method of analysis (ASBC, 2011). The control strains produced expected flocculence values: for example, the Belgian strain (WLP570) is non-flocculant (1.9%) and the British strain (WLP002) is highly flocculant (98.3%) (Table 3). We observed high levels of flocculation among the *kveik* yeasts, but this property was surprisingly not universal: 16/24 strains produced flocculence values >70%, while others showed very low flocculence (<20%). Interestingly, in most *kveik* samples, except for Muri and Joniskelis, at least one of the isolated strains showed high flocculation rates above 88% (Table 3). It is possible that in the original *kveik* mixed *S. cerevisiae* cultures, the yeasts undergo co-flocculation and consequently some strains never developed this function (Rossouw, Bagheri, Setati, & Bauer, 2015; Smukalla et al., 2008). Nonetheless, the high incidence of efficient flocculation among *kveik* yeasts is further support these yeasts have been domesticated, and may contain the copy number variations linked to flocculation genes (*FLO*) which are common among domesticated yeasts (Bergström et al., 2014; Dunn, Richter, Kvitek, Pugh, & Sherlock, 2012; Gallone et al., 2016; Steenwyk & Rokas, 2017)

## Discussion

Here we present evidence that suggest *kveik* yeasts obtained from Norwegian farmhouse brewers represent a previously undiscovered branch of the beer yeast family tree (Almeida et al., 2015; Baker et al., 2015; Gallone et al., 2016; Gonçalves et al., 2016), and that these yeasts have promising beer production attributes. Our PCR fingerprint data, assembled with two interdelta primer sets, accurately resolved similar groups among known English, American and German ale strains, and clearly showed individual *kveik* yeast strains form a genetically distinct group of ale yeasts (Fig. 2). Moreover, our analysis suggests these yeasts cluster genetically with geographic provenance separated by the Jostedal glacier (Fig. 1, 2).

Our investigation of the beer production attributes with small-scale fermentation trials and phenotypic screens revealed the majority of the Norwegian *kveik* yeasts metabolize wort sugars quickly, are POF-, flocculate efficiently, are highly ethanol tolerant and thermotolerant (Fig 3 & Table 3). Previous genetic analyses of known beer strains attributed such characteristics of domestication to specific genetic determinants, and some combinations of these determinants are typically present in industrial beer yeasts (Gallone et al., 2016; Gonçalves et al., 2016). However, it is rare to find all these domestication phenotypes combined in a single yeast strain as is the case for several of the *kveik* strains. The increased production rates of early industrial breweries in the 17^th^-18^th^ century was previously proposed to provide the foundation for beer yeast domestication (Gallone et al., 2016). Here we show *kveik* yeasts, surprisingly, have similar adaptation characteristics to the beer fermentation environment despite being domesticated by farmhouse brewers without the high-frequency production pressure of an industrial brewing environment (Gallone et al., 2016). In combination, our observations suggest that either: *kveik* yeasts were domesticated independently of, and potentially earlier than other known domesticated beer strains; the process of yeast domestication can occur on a generational scale shorter than previously hypothesized; or that specific aspects of the traditional *kveik* fermentation process can accelerate domestication.

Several phenotypes of domestication have been attributed to different genetic determinants. For example, maltotriose fermentation is critical to utility of yeasts for beer production, but is not a common feature among wild yeast populations (Gallone et al., 2018). This trait has evolved through two genetic pathways among beer yeasts. In one case, a mutation in *AGT1* (a high-affinity maltose transporter) is present conferring an increased affinity for maltotriose (Alves et al., 2008). The second group of yeasts contain a non-functional *AGT1* allele, indicating that a presently unknown mechanism mediates maltotriose fermentation in these yeasts, with some evidence that this is a polygenic trait (Brown et al., 2010). It is also likely that the ubiquity of POF-yeasts in brewing is due to a human preference for these yeasts, and that POF-yeasts, containing mutations in *PAD1* and *FDC1* consequently have become widespread among brewing yeasts (Gallone et al., 2016, 2018; McMurrough et al., 1996). Given our results, we expect *kveik* yeasts to show one or more adaptations conferring enhanced maltotriose fermentation as well as loss-of-function mutations in *PAD1* and/or *FDC1* conferring POF-.

Also, flocculation in yeast is driven by *FLO* genes, which encode different cell-surface adhesins that enable neighboring cells to clump together to form visible flocs that settle out of suspension (Smukalla et al., 2008). In beer yeasts, many of these genes have copy number variations (CNVs) which may serve to enhance flocculation (Gallone et al., 2016; Steenwyk & Rokas, 2017). Similarly, deletion of *SAS2*, an acetyltransferase, also result in enhanced flocculation and maltose fermentation (Rodriguez, Orozco, Cantoral, Matallana, & Aranda, 2014). While the genomic adaptations in *kveik* yeasts are currently unknown, it is feasible to suggest that CNVs and gene deletions, as mentioned for *FLO* genes and *SAS2*, respectively, may represent potential genomic alterations that would explain both the enhanced flocculation and fermentation efficiency of *kveik* yeasts. Approximately one third of the *kveik* yeasts did not flocculate with high efficiency. This may be influenced by the procedure used by farmhouse brewers to harvest yeast for repitching, including harvesting at least some of the top-fermenting yeast cells where the evolutionary pressure to flocculate would be less. It is therefore not surprising that some kveik strains flocculate less efficiently than others.

Wort fermentations revealed that *kveik* strains produce a range of fruity esters, with ethyl caproate, ethyl caprylate, ethyl decanoate, and phenethyl acetate present above detection threshold (Table 3), indicating that these yeasts can be used to produce beers with fruity character. How *kveik* yeasts compare to a broader range of industrial beer yeasts in terms of diversity and intensity of flavour production is currently unknown. In addition, we have shown the *kveik* ale yeasts have a broad range of wort attenuation values (Fig. 3). As these yeasts are POF-, a desirable trait for the majority of beer styles (McMurrough et al., 1996), they also could have broad utility as a new group of ale yeasts, with selection by the brewer in accordance with desired attenuation target values and flavour profiles.

Strikingly, our phenotypic screening revealed the enhanced thermotolerance and ethanol tolerance of these yeasts in comparison to known domesticated beer yeasts (Table 3). Long-term heat adaptation is particularly relevant to fermentation processes performed at elevated temperatures, including those used for industrial bioethanol production. Multiple molecular and cellular processes and targets have been identified in the adaptation of yeast to heat. A prior study investigating the adaptation of yeast to ~40 °C over a prolonged period of time, identified SNVs (single nucleotide variations) in genes related to DNA repair, replication, membrane composition and membrane structure as specific genetic markers of thermotolerance (Caspeta et al., 2013). Furthermore, large genetic duplications were identified in thermotolerant yeasts, in line with other evidence for gene duplication as a typical mechanism for adaptation in *S. cerevisiae* (Caspeta et al., 2013; Gallone et al., 2016). Moreover, the study identified nonsense mutations in *ERG3*, a C-5 sterol desaturase, which resulted in an altered sterol composition of the plasma membrane rendering the yeast more thermotolerant.

Interestingly, high-temperature exposure also renders yeast more amenable to desiccation (Welch, Gibney, Botstein, & Koshland, 2013), a trait presumably common among *kveik* yeasts given that many traditional brewers dry their yeast [ref]. Deletion of *SCH9*, a signaling kinase that function in controlling cellular adaptation, increases the desiccation tolerance of yeast cells, and can also accelerate growth rate (Welch et al., 2013). Although the mechanism(s) of thermotolerance in *kveik* yeasts are currently uncharacterized, genomic alterations such as SNVs/SNPs and genomic duplications selected for with repeated exposure to high-temperature and desiccation environments, can help explain the high temperature and presumed desiccation tolerance of *kveik* yeasts. The thermotolerance characteristic has potential application in brewing as wort inoculation at higher temperatures (>30 °C) without compromise in flavour could help limit the expensive cooling needed to manage wort fermentation temperatures that are typically controlled at 18-22 °C for ale fermentations (Hill, 2015).

We also demonstrate that ethanol tolerance is a common adaptation of *kveik* yeasts. Ethanol tolerance is known to be a polygenic and genetically complex trait involving multiple alleles. While single genetic alterations can incrementally increase ethanol tolerance, it does not approach that of the polygenic/multiallelic phenotype (Lam, Ghaderi, Fink, & Stephanopoulos, 2014; Snoek, Verstrepen, & Voordeckers, 2016). High ethanol environments generally disrupt cell membrane structure and function, and impact protein folding. Not surprisingly, genes linked to ethanol tolerance are often associated with: stabilizing cell walls and cell membranes; increasing the protein folding capacity; maintaining the electrochemical gradient across the plasma membrane; and maintaining vacuolar function to mention a few (Lam et al., 2014; Snoek et al., 2016). Remarkably, almost one third of the *kveik* yeasts reported here could grow in the presence of 16% ethanol. Given the ethanol and high temperature tolerances of *kveik* yeasts, it is possible these yeasts could benefit the distillation and bioethanol industries where these traits are desired (Caspeta et al., 2016).

Whole genome sequencing has previously been used to delineate groups of industrial yeasts genetically (Almeida et al., 2015; Baker et al., 2015; Gallone et al., 2016; Gonçalves et al., 2016). Such an approach would provided increased resolution not only of the positioning of the *kveik* yeasts in the domesticated beer yeast tree, but also allow for the direct genomic comparison with the now extensive range of industrial yeast whole genome assemblies available (Almeida et al., 2015; Gallone et al., 2016; Gonçalves et al., 2016). Perhaps most importantly, a whole genome sequencing approach would provide insight into the specific changes in domestication markers which have occurred in the *kveik* yeasts, and demonstrate whether these are identical or different to the mutations which have occurred in other domesticated yeasts (Brown et al., 2010; Gallone et al., 2016; Gonçalves et al., 2016; McMurrough et al., 1996; Steensels & Verstrepen, 2014). To our knowledge, the genomic sequences of kveik yeasts have not been determined and will be pivotal in understanding the domestication and characteristics of these yeasts.

## Acknowledgements

We would like to acknowledge Lars Marius Garshol for his assistance in obtaining yeast cultures, as well as critical review of the manuscript. We also thank Cam Fryer and Royal City Brewing for the donation and production of wort used in this study This research was funded by an NSERC Discovery (#264792-400922) and OMAFRA-University of Guelph Gryphons LAAIR (LAAIR2015-5142) grants.

